# Occurrence of Avian Reovirus and Picobirnavirus in Wild Birds in an Environmental Protection Area in the Amazon Biome, Pará, Brazil

**DOI:** 10.1101/2022.01.11.475455

**Authors:** Diego Pereira, Lizandra Caroline dos Santos Souto, Sylvia de Fátima dos Santos Guerra, Edvaldo Tavares da Penha Júnior, Patrícia dos Santos Lobo, Luana da Silva Soares, Helder Henrique Costa Pinheiro, Elaine Hellen Nunes Chagas, Bruna Alves Ramos, Liliane Leal das Chagas, Maria Nazaré Oliveira Freitas, Erilene Cristina da Silva Furtado, Jéssica Cecília Pinheiro Rodrigues, Alexandre do Rosário Casseb, Lívia Caricio Martins, Joana D’Arc Pereira Mascarenhas

## Abstract

Wild birds have great prominence on transmission of diseases to humans, mainly due to their ease of access to human population, raising concerns about the potential impact of that proximity in context of the One Health. Studies referring to circulation of avian reovirus (ARV) and picobirnavirus (PBV) in wild birds are limited, in addition to reinforcing the development of researches that describe the prevalence, characterize the variants and evaluate the potential impact of these infections on the wild ecosystem and public health. The present study reports the occurrence of ARV and PBV in wild birds collected from an environmental protection area in the Amazon biome. RT-PCR analysis showed ARV infection prevalence in 0.6% (1/155) and PBV infection in 1.29% (2/155) on the samples. ARV strain isolated in this study demonstrated more phylogenetically related to other ARVs previously circulating in poultry in the same region. The two PBV strains obtained belong to genogroup I, and showed phylogenetically related to other PBV isolated from different animal species in different geographic regions. This study is a pioneer in the detection of ARV in wild birds in Brazil and presents a report of the first occurrence of PBV in wild birds of *Guira guira* specie. Additional studies in wild birds are required to increase the epidemiology, origin, evolution and emergence of new viruses that may provoke problems in the context of One Health.

## INTRODUCTION

Wild birds are among the animals that have great prominence in context of disease transmission to humans. Over the years, these animals approached even closer to humans being, due to beautiful plumage, songs and colors^1^. The ability to fly propitiate to birds an ease of access in places close to the human population, however these environments became a local of high risk to transmission of pathogens to humans, raising concerns about potential impact of this proximity in context of One Health^2^.

Viruses are among the most important clinical and epidemiological pathogens in birds, as infections that occur in the first weeks of life in avian species are usually of viral etiology^3,4,5^. Rotaviruses (RV), Avian reoviruses (ARV), Picobirnaviruses (PBV), Avian Influenza viruses (AIV), Astroviruses (AstV), Coronaviruses (CoV) and West Nile viruses (WNV) are examples of the most important viruses for global public health. This is due to the potential for dispersal of wild birds, especially those that have migratory routes^6,7,8,9,10,11,12,13^.

ARVs and PBV are frequently reported in infecting poultry, associated with clinical or subclinical disease, causing seriously economic impacts to poultry industry^14,15,16,17^. ARVs are described as important agents that provoke gastroenteric diseases, viral arthritis and tenosynovitis, in birds^18,19^. On the other hand, PBV may be detected in normal or diarrheal excrement from domestic and wild birds, therefore its role as a primary agent of acute gastroenteritis remain unestablished, reinforcing the importance of studies that had better characterize the pathogenic aspects of this microorganism. ^20,21^.

ARVs belong to the family *Reoviridae*, subfamily *Spinareovirinae* and genus *Orthoreovirus*^22^. Its capsid has 70-85 nm, icosahedral symmetry and it has no lipoprotein envelope^23^. The genome is composed of ten double-stranded RNA segments (dsRNA)^24^. PBVs belong to the family *Picobirnaviridae* and the genus *Picobirnavirus*^25^. The viral particle contains around 33-41 nm, icosahedral symmetry and presents no envelope^26^. Its genome consists of bi-segmented dsRNA, where segment 2 classifies PBVs into genogroup I (GI), genogroup II (GII), and non-I and non-II genogroup^27^.

In Brazil, there is a lack of research referring to the occurrence of ARV and PBV in free-living wild birds. Especially species that live in environmental protection areas close to large urban centers, where there is a great risk of zoonotic transmission of infectious agents due to the large variety of wild animals present, and their proximity to the human population. Molecular epidemiology studies are essential to describe the prevalence of infectious agents, characterizing the variants present and estimate the potential impact of infections on the wild ecosystem and in public health.

In this context, due to the limited studies on the circulation of these viruses in wild birds, this research described the occurrence of ARV and PBV in wild birds collected from an environmental protection area in the Amazon biome.

## METHODS

### Study area

The study area included forest areas close to deforested areas for grassland and/or for buildings construction, in the territory belonging to Federal Rural University of Amazon (UFRA) (1° 27’ 21” S 48° 26’ 12” W). The university UFRA is located in the Environmental Protection Area of Metropolitan Region of Belém (APA - Metropolitan Belem), which presents a total protected area of 5.647 hectares (ha) and 56.47 square kilometers (km^2^). APA (Metropolitana Belém) is an environmental conservation unit of the Pará state for sustainable use located in the Amazon biome. This environmental conservation area admit a wide variety of wild animals in preservation, thence human activity is limited.

### Bird capture and collection of clinical specimens

From February to October 2019, birds were collected in three distinct climatic periods (rainy, dry and intermediate). To the capture, were using mist nets fixed to the ground with metal wattle, stretched from 5:00 am to 10:00 am, and checked every thirty minutes. After capture, took note some information as weight and taxonomy based on morphological aspects (order, family and species), genus (male or female) and life stage (young or adult)^28,29,30^.

The birds were kept individually in cardboard boxes lined with aluminum foil paper. The fecal specimens were collected of the box or through the gentle introduction of sterile swab directly into the cloaca, placed in cryogenic tubes and stored at −20°C until processing. After collecting the specimens, the birds were marked with non-toxic ink (Raidex®) to identification in case of recaptured and after released back into the environment. Due to stress, some birds died during the process of capture, wherefore it was not possible to collect the feces, the species were submitted to necropsy and the intestine samples were stored.

### Viral genome extraction

Suspensions were prepared at 10% by diluting feces and/or intestinal contents in Tris/HCl/CaCl^2+^ buffer (pH 7.2 0.01M), clarified by centrifugation at 4.000 rpm/10 minutes. The supernatant was submitted to viral genome extraction according to the protocol described by Boom et al.^31^.

### Polyacrylamide Gel Electrophoresis (PAGE)

The products of extraction were submitted to PAGE for detection of ARV and PBV by electrophoretic profiles according to the technique described by Pereira et al.^32^.

### RT-PCR for ARV

RT-PCR was performed targeting ARV S2 gene. To amplify a partial fragment of 625bp of the S2 gene, the forward primer PAF (5’ - ACT TCT TYT CTA CGC CTT TCG - 3’) and the reverse PAR (5’ - ATY AAW DCW CGC ATC TGC TG - 3’) were used^33^. To obtain the complementary DNA strain (cDNA), 4 μL of extracted dsRNA and 2 μL of pair of primers (20 mM) were used. The reaction followed an incubation of 5 minutes at 97°C for denaturation of the dfRNA, followed by 5 minutes at 0°C for heat shock.

Reverse transcription was performed to a final volume of 25 μL. This mix was obtained by adding 19 μL of RT mixture including 11 μL of DNAse/RNAse free H_2_O (Hyclone^™^), 1 μL of dNTPs (20mM, Promega^®^), 5 μL of buffer (5x, Promega^®^), 1.5 μL of MgCl_2_ (25 mM, Promega^®^) and 0.5 μL of RT (4U, Promega^®^), followed by an incubation at 42°C for 60 minutes. After reverse transcription, PCR was performed, adding to the cDNA 25 μL of the PCR mixture containing 15.25 μL of H_2_O free DNAse and RNAse (Hyclone^™^), 3 μL of dNTPs (20mM, Promega^®^), 5 μL of buffer (5x, Promega^®^), 1.5 μl of MgCl_2_ (25mM, Promega^®^) and 0.25 μl of Taq DNA Polymerase (5U, Promega^®^). The cycling conditions used were described by Zhang et al.^33^.

The amplicons obtained by PCR were performed using agarose gel electrophoresis, concentration of 1.5% in Tris/Borate/EDTA (TBE) buffer and gel stained with SYBR^®^ Safe DNA Gel Stain (Invitrogen^®^). GEL DOC 1000 image processor (Bio-Rad Laboratories, Inc., Hercules, CA) performed photo documentation.

### RT-PCR for PBV

RT-PCR was performed targeting the PBV RdRp gene. To amplify a 201bp genogroup I fragment, PicoB25 forward primers (5’-GCN TGG GTT AGC ATG GA-3’) and PicoB43 reverse (5’-A(GA)T G(CT)T GGT CGA ACT T-3’)) were used^34^. For genogroup II, PicoB23 forward primers (5’-CGG TAT GGA TGT TTC-3’) and PicoB24 reverse (5’-AAG CGA GCC CAT GTA-3’) were used to amplify fragments of 369bp^34^. To obtain the cDNA, 4 μL of extracted dsRNA and 1 μL of primer pair (20mM) were used, followed by 5 minutes incubation at 97°C for dsRNA denaturation, and 5 minutes of heat shock at 0°C.

To the first step, reverse transcription, followed denaturation were added 20 μL of the RT mixture containing 12.25 μL of DNAse/RNAse free H_2_O (Hyclone^™^), 1 μL of dNTPs (20mM, Promega^®^), 5 μL of buffer (5x, Promega^®^), 1.5 μL of MgCl_2_ (25mM, Promega^®^) and 0.25 μL of RT (4U, Promega^®^), followed by an incubation at 42°C for 60 minutes. The second step, PCR, were added to cDNA 25 μL of the PCR mixture containing 15.25 μL of DNAse/RNAse free H_2_O (Hyclone^™^), 3 μL of dNTPs (20mM, Promega^®^), 5 μL of buffer (5x, Promega^®^), 1.5 μL of MgCl_2_ (25mM, Promega^®^) and 0.25 μL of Taq DNA Polymerase (5U, Promega^®^). The cycling conditions used were described by Silva et al.^20^.

The amplicons obtained by PCR were performed using agarose gel electrophoresis, concentration of 1.5% in Tris/Borate/EDTA (TBE) buffer and gel stained with SYBR^®^ Safe DNA Gel Stain (Invitrogen^®^). GEL DOC 1000 image processor (Bio-Rad Laboratories, Inc., Hercules, CA) performed photo documentation.

### Nested-PCR

Samples that presented amplicons of 201bp in the previous RT-PCR for PBV GI were submitted to a new RT-PCR followed by a Nested-PCR, to amplify a larger region of the RdRp gene. The forward primers PBV 1.2F (5’-AAG GTC GGK CCR ATGT-3’) and reverse PBV 1.2R (5’-TTA TCC CYT TTC ATG CA-3’) were used to amplify a fragment of 1229bp^35^. In Nested-PCR, the Malik-2-FP forward primer (5’-TGG GWT GGC GWG GAC ARG ARGG-3’) and the Malik-2-RP reverse (5’-YSC AYT ACA TCC TCC AC-3’) were used, which amplify a fragment of 580bp of RdRp gene^35^. The cycling conditions used were those described by Malik et al.^35^.

### Nucleotide sequencing and phylogenetic analysis

Amplicons were purified using the ExoSAP-IT^™^ kit (Applied Biosystems^™^) according to the manufacturer’s recommendations. Next purification, the products were subjected to nucleotide sequencing using RT-PCR/Nested-PCR primers, and the Big Dye Terminator^®^ v.3.1 kit (Applied Biosystems^™^) according to the manufacturer’s recommendations. Final reaction was submitted to ABI PRISM 3130 Automated Genetic Sequencer (Applied Biosystems^™^).

The sequences were edited using BioEdit v.7.2 program, aligned by MEGA v.10.0.537 program^36^ and compared with other sequences deposited in GenBank (www.ncbi.nlm.nhi.gov) through the Basic Local Alignment Search Tool (BLAST)^37^. Phylogenetic trees were constructed by MEGA v.10.0.537 program^36^ using Neighbor-Joining method and Kimura two-parameter model^38^. Bootstrap of 2000 replicates was used to phylogenetic groups^39^. Nucleotide similarities were calculated by Geneious v.10.0.7 program^40^.

### Statistical analysis

Data analysis was executed using BioEstat v.5.3 program^41^. G test and Fisher’s exact statistical test were performed to verify differences in the results of PBV and ARV occurrence between the categorical variables of this study. Differences were considered significant at 5%.

## RESULTS

A total of 155 clinical specimens from wild free-living birds were collected from February to October 2019. The fecal specimens were 144 and intestine specimens were 11. The species belong as following orders: Passeriformes (n=127), Columbiformes (n=9), Cuculiformes (n=8), Psittaciformes (n=3), Coraciiformes (n=3), Caprimulgiformes (n=3), Apodiformes (n=1) and Accipitriformes (n=1). All samples were tested by PAGE and RT-PCR for ARV and PBV.

The PAGE test showed negativity for all of the specimens tested. The result presented no Electrophoretic profile consistent with ARV and PBV, except the positive controls used during technique.

Referring to RT-PCR for ARV S2 gene, 0.6% (1/155) of positivity was observed in a specimen from a wild bird long-winged-antwren (*Myrmotherula longipennis*). For PBV GI 1.29% (2/155) were positives, from wild bird guira-cuckoo (*Guira guira*). Therefore, there was no positive samples for PBV GII.

Regarding to ARV and PBV positivity among the different study variables, the statistical analysis did not show significant differences (**Table 1**).

**Table 1.** Statistical analysis of the frequency of ARV and PBV positivity in wild birds according to the different variables of this study. ND - Not defined; p - Probability value calculated by statistical test; *Test G; # Exact test of Fisher.

| Variable | PBV |  |  |  | p | ARV |  |  |  | p |
| --- | --- | --- | --- | --- | --- | --- | --- | --- | --- | --- |
|  | Positive |  | Negative |  |  | Positive |  | Negative |  |  |
|  | n | % | n | % |  | n | % | n | % |  |
| Sex |  |  |  |  | 0,383* |  |  |  |  | 0,320* |
| Female | 0 | 0,0 | 26 | 100,0 |  | 0 | 0,0 | 26 | 100,0 |  |
| Male | 0 | 0,0 | 50 | 100,0 |  | 1 | 2,0 | 49 | 98,0 |  |
| ND | 2 | 2,5 | 77 | 97,5 |  | 0 | 0,0 | 79 | 100,0 |  |
| Age group |  |  |  |  | 0,980* |  |  |  |  | 0,922* |
| Young | 0 | 0,0 | 11 | 100,0 |  | 0 | 0,0 | 11 | 100,0 |  |
| Adult | 2 | 1,4 | 141 | 98,6 |  | 1 | 0,7 | 142 | 99,3 |  |
| ND | 0 | 0,0 | 1 | 100,0 |  | 0 | 0,0 | 1 | 100,0 |  |
| Sample type |  |  |  |  | 1,000<sup>#</sup> |  |  |  |  | 0,405<sup>#</sup> |
| Feces | 2 | 1,4 | 142 | 98,6 |  | 1 | 0,7 | 143 | 99,3 |  |
| Intestine | 0 | 0,0 | 11 | 100,0 |  | 0 | 0,0 | 11 | 100,0 |  |
| Climatic period |  |  |  |  | 0,168* |  |  |  |  | 0,301* |
| Rainy | 0 | 0,0 | 39 | 100,0 |  | 0 | 0,0 | 39 | 100,0 |  |
| Dry | 0 | 0,0 | 69 | 100,0 |  | 0 | 0,0 | 69 | 100,0 |  |
| Intermediary | 2 | 4,3 | 45 | 95,7 |  | 1 | 2,1 | 46 | 97,9 |  |

The partial sequences of the PBV RdRp gene of the two strains obtained in this study were compared with other prototype PBV sequences isolated in Brazil and in other countries and deposited in GenBank. Phylogenetic analysis grouped two strains isolated in this study into PBV GI, however, the two sequences were heterogeneously related, grouping divergently in the tree (**Fig. 1**).

**Fig. 1.**
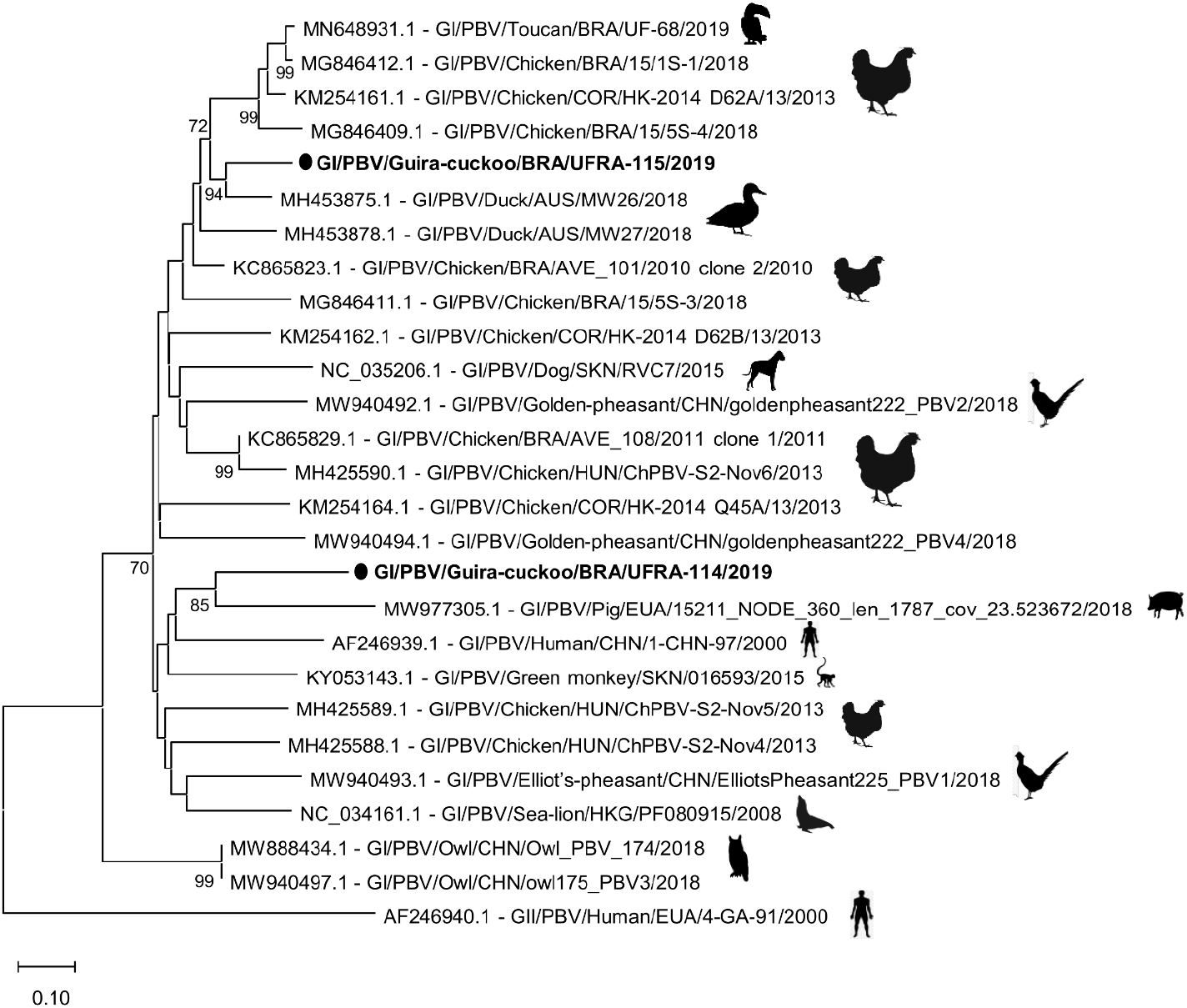
Phylogenetic tree based on partial sequence alignment of PBV RdRp gene. The sequences of this study are represented in bold. The numbers next to the nodes indicate boostrap values >70%. The scale bar is proportional to phylogenetic distance. The prototype strain GII/PBV/Human/USA/4-GA-91/2000 (AF246940.1) was used as an external group to better understand the phylogenetic relationships between the strains. The phylogenetic tree was constructed using Neighbor-Joining method and Kimura two-parameter model, with bootstrap of 2000 replicas to give consistency to the phylogenetic groups.

The strain GI/PBV/Guira-cuckoo/BRA/UFRA-115/2019 grouped with a sequence from a duck in Australia (MH453875.1), exhibiting a bootstrap of 94%. This grouping was phylogenetically related to other strains isolated from toucan and chickens in Brazil and South Korea (bootstrap of 72%).

The strain GI/PBV/Guira-cuckoo/BRA/UFRA-114/2019 grouped with a strain from swine in the USA (MW977305.1), presenting bootstrap of 85%. This cluster presented phylogenetically related to other PBV isolated from mammals.

The homology between PBV sequences of this study represented 59.5% of nucleotide identity, showing a high genetic diversity. When compared with prototype of PBV sequences, the values were from 55.0 to 81.4%. The GI/PBV/Guira-cuckoo/BRA/UFRA-115/2019 strain showed higher nucleotide similarity (81.4%) with a sequence obtained from duck reported in Australia in 2018 (MH453875.1), and two isolated prototypes from chickens from Brazil and South Korea (KC865823.1 and KM254161.1), with 76.6 and 75.6%, respectively.

The PBV strain (GI/PBV/Guira-cuckoo/BRA/UFRA-114/2019) showed higher nucleotide similarity, 63.4, 61.9 and 60.7%, with two sequences obtained from chickens in Brazil (KC865823.1 and KC865829.1), and a prototype obtained from a green monkey (KY053143.1) reported in 2015 on Saint Kitts and Nevis, an island in the Caribbean.

The partial sequence of ARV S2 gene of the present study was compared with other ARV prototype sequences obtained from domestic birds (chicken, turkey, duck and goose) and wild birds (pheasant, wild duck, brown-eared bulbul and crow) isolated in Brazil and other countries around the world. The REO/Long-winged-antwren/BRA/UFRA-118/2019 strain was phylogenetically related to ARV strains previously reported in chickens in Brazil (KY783741.1, KY783739.1), with a bootstrap of 85% (**Fig. 2**).

**Fig. 2.**
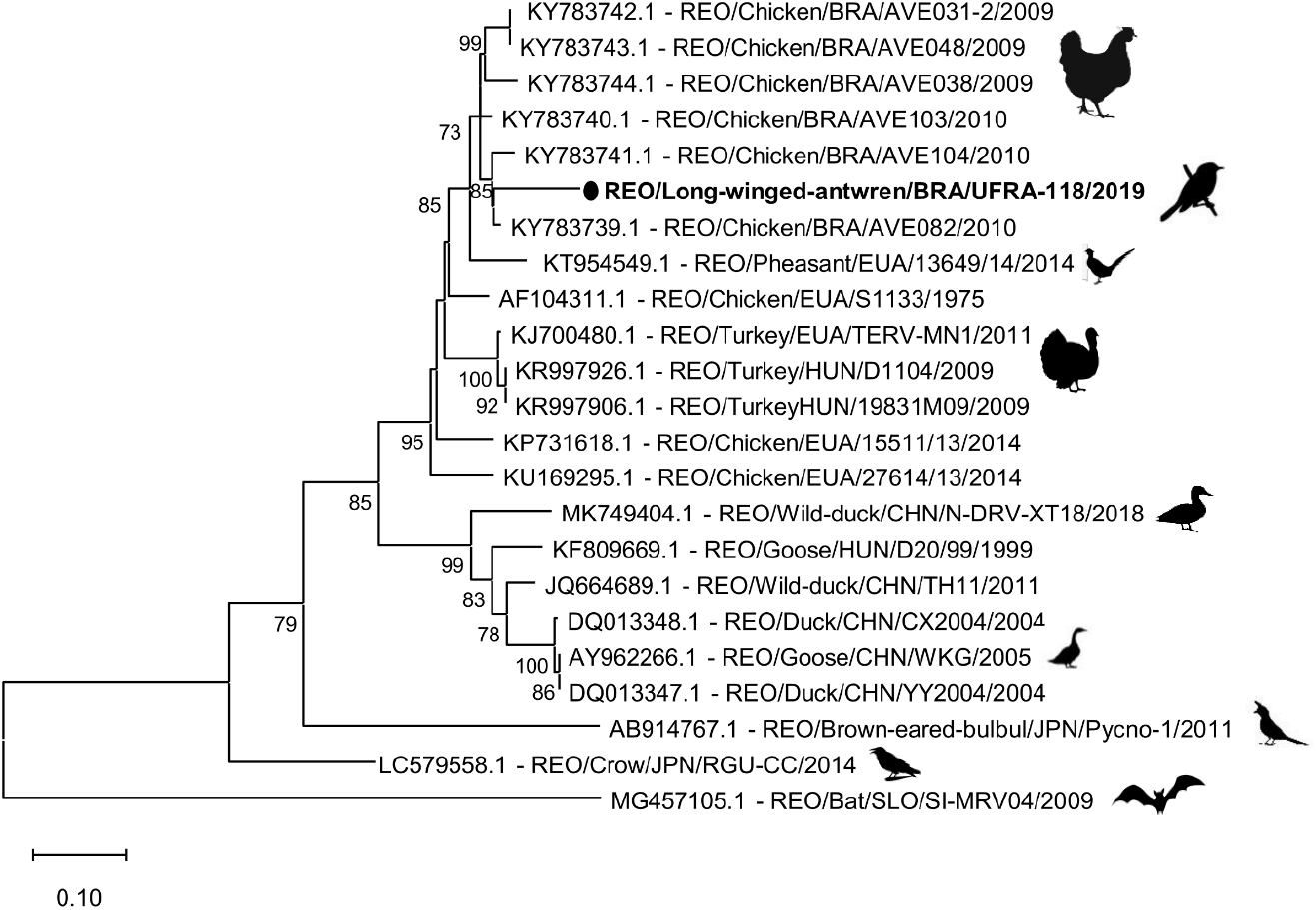
Phylogenetic tree based on the alignment of partial sequences of the ARV S2 gene. The sequence of the present study is represented in bold. The numbers next to the nodes indicate boostrap values >70%. The scale bar is proportional to phylogenetic distance. The strain REO/Bat/SLO/SI-MRV04/2009 (MG457105.1) represents a prototype of MRV and was used as an outgroup to better understand the phylogenetic relationships between the strains. The phylogenetic tree was constructed using Neighbor-Joining method and Kimura two-parameter model, with bootstrap of 2000 replicas to give consistency to the phylogenetic groups.

Regarding to nucleotide identity, S2 gene sequence, in this study, showed from 51.9% to 86.4% similarity when comparing with prototype ARV sequences used. The highest homologies observed were sequences obtained from chickens in Brazil (KY783741.1, KY783743.1, KY783742.1) collected in 2009 and 2010, with 86.4, 86.0 and 85.8% of similarity. However, the lowest homology was a sequence obtained from brown-eared bulbul (AB914767.1) isolated in Japan in 2011, with 51.9% of similarity.

## DISCUSSION

ARVs and PBV are among enteric important viruses that have been widely reported infecting poultry and wild birds, producing noted impacts to poultry economy and wildlife preservation^7,12,19,20,21,42^. The characteristic of segmented genome of ARV and PBV is an important factor that contributed to rapid dispersion, evolution and adaptation of these viruses to different hosts. Therefore, investigation is needed, especially in wild birds, which are known to be important reservoirs of several infectious agents^26,43,44,45,46^.

In the present study, there was no electrophoretic profiles to ARV and PBV in PAGE migration, corroborating other previous studies involving wild birds specimens in Northern Brazil, that showed the same result^12,47,48^. However, previous studies in Brazil and other countries have already characterized electrophoretic profiles of PBV and ARV in poultry (*Gallus gallus domesticus*), domestic ducks (*Cairina moschata*) and wild birds (*Hypsipetes amaurotis*)^20,44,49^.

Several studies have suggested that low frequency of detection of ARV and PBV by PAGE is associated with limiting factors such as sample quality, time elapsed from collection to analysis, low viral load in animal feces and host factors such as age and immunological system^26,50,51^. Although PAGE exhibits limited sensitivity, this technique has been highly specific, due to the characteristic electropherotype of each virus, several genomic variations contribute to the difficult detection by primers in RT-PCR^26,52^.

The prevalence of PBV by RT-PCR in the present study reported 1.29% of positivity. Studies carried out in Brazil that investigated the presence of PBV in wild birds found positivity ranging of 0 - 4.5 %, using the same technique and target gene^12,47,48^. Higher prevalence of PBV infection were detected in studies with poultries, reported the frequency of 11.76 - 49.41% of positivity in broilers (*Gallus gallus domesticus*), and 51.67% in turkeys (*Meleagris gallopavo*)^20,21,51^.

The high positivity of PBV in studies with broilers, especially on poultry farming, suggest that positivity may be associated to confined system these birds are raised in most of the time, facilitating viral dispersion between them, due to direct contact with each other^52^. Due to this study works with free-living wild birds, the probability of virus spread among the species was reduced, even though some avian species that live in wild environments presents habit of living in flocks.

The low occurrence of PBV was attributed to limitation of primers used in RT-PCR, which were designed for PBV of human origin^34^. Several studies demonstrated high sensitivity that initiators in detecting these microorganism and the genogroup in different hosts^35,53^. On the hand, some studies indicated a limited effectiveness for these initiators^26^. Another important point to highlight is the existence of other PBVs belonging to different genogroup, which was not detected by initiators on this study, underestimating the real prevalence of PBV in the analyzed specimens^48^.

The genogroups GI and GII were investigated in wild birds in this study, however GI was detected in 1.29% of the specimens, corroborating other previous studies^12,47,54^. Chagas^47^ detected GI in wedge-billed-woodcreeper (*Glyphorynchus spirurus*) captured from a deforested area in Santa Bárbara city, Pará, Brazil. Duarte Júnior et al.^12^ detected GI in a sample from toucan (*Rhamphastus sp*.) with diarrhea symptoms, and treated in a veterinary hospital in Castanhal city, Pará, Brazil. In Australia, a study characterized three PBV sequences of GI from fecal specimens from Australian duck (*Tadorna tadornoides*)^54^.

The prevalence of ARV in wild birds in the present study was 0.6% (1/155), showing divergence when compared with other previous studies^55,56^. In Brazil, only two studies investigated ARV S2 gene in wild bird fecal specimens, however no positivity was reported^47,48^. Silva^52^ described positivity in 32.9% (28/85) in poultry fecal specimens raising on poultry farming, for the ARV S2 gene. In 2019, a report in Egypt found ARV prevalence of 33.3% (5/15) in poultry, to S2 gene as a target^56^. This study is the first report regarding to occurrence of ARV in wild free-ranging birds in Brazilian territory.

In studies that reported high prevalence of ARV infection, specimens come from birds with clinical signs of infection, such as arthritis, tenosynovitis, enteric syndromes, in addition to more severe cases when central nervous system is compromised and death^24,55^. In USA, Lu et al.^7^ investigated ARV from tendons, synovial tissues and viscera from chickens, turkeys, partridges, quails and pheasants, that presented clinical signs, and all of 311 specimens analyzed on that study were positive for ARV. In the present study, no clinical signs information was collected, therefore this aspect was not considered on discussion.

Reports in regard to occurrence of ARV in domestic and wild birds seem limited, focusing on describing molecular epidemiology in birds, experimental aspects of ARV infection in animal models, viral isolation in cell culture, and complete genome sequencing of specific strains^8,42,44,57,58^. Therefore, the present study contribute to understanding molecular epidemiology of ARV in free-living wild birds.

Studies related to PBV described that strains isolated in geographically distinct locations and from different hosts presented greater phylogenetic relationship between them, when compared with strains isolated in the same study and in the same region^20,21,35,54,59^. The findings of this study corroborate the literature in relation to rapid spread, evolution and adaptation of PBV, which suggest that the great similarity between lineages belonging to different geographic regions may be related to sharing of the same ancestor and also reduction of trade barriers between countries^26,51^.

PBV strains isolated in this study were collected from wild birds belong the same species (*Guira guira*), captured on the same day and location, and the great genetic diversity observed corroborates heterogeneous nature of PBVs described on literature. Several factors are considered responsible of high genetic diversity of PBVs, such as small size of the analyzed fragment, genetic variability, multiple interspecies transmissions and genetic rearrangement events between segments of different PBV strains^20,26,51^.

PBV GI shows a worldwide distribution pattern and has been reported to infect a variety of hosts such as mammals, birds, reptiles, and even fish^60,61,62,63^. The phylogenetic grouping of PBV GI isolated from different hosts indicates these viruses are not species specific, although, they can transmit from one host to another^21,35,47^. The great genetic similarity observed between PBV strains isolated from different hosts raises increasing concerns about the zoonotic potential of this virus in the context of One Health^26,64^.

Silva^52^, in studies with S2 gene in poultry, showed that phylogenetic analysis in 13 of 15 ARV strains isolated reported more phylogenetically related to each other, while two of them showed more divergence in the tree, and were closer to the ARV prototypes of chicken, turkey and ostrich used in comparison. All of 15 strains showed a nucleotide homology of 90.1-100% between them, and 90.9-94.4% with the reference prototypes achieved from other production birds around the world, demonstrating the high degree of nucleotide similarity that strains of ARVs obtained in the same location presented among them, and with strains isolated from different geographic regions.

The high nucleotide identity and phylogenetic inference showed that ARV strain isolated from the wild bird long-winged-antwren (*Myrmotherula longipennis*) in the present study was more related to ARV strains circulated for some years in poultry on the same region than strains circulated in other countries of the world. These findings suggest that ARV strains circulating in the same region may be adapted to the environment over the years, and ARV transmission between domestic and wild birds may occur on nature, as commercial poultry farming normally localized in rural environments close to wild environments, favoring close contact between these animals and the transmission of pathogens.

The occurrence of ARV and PBV in wild birds corroborated studies that also identified the circulation of these viruses in population of birds. The findings reported here suggest that the wild species of birds: *Guira guira* and *Myrmotherula longipennis* may act as reservoirs of PBV and ARV infection, respectively. Furthermore, even these birds are not considered migratory, they may still act as possible dispersion agents for these viruses in wild and urban environments.

The wild birds were collected in an Environmental Protection Area (APA), which consists of an extensive forest fragment with limited human activity and wildlife preservation. The circulation of ARV and PBV in this environment demonstrated the cycles of these agents occur naturally in the wild ecosystem and the fact that APA – Metropolitana Belém be formed by several populated neighborhoods and poor basic sanitation infrastructure, possible events transmission between avian species and humans can trigger.

In conclusion, this study is a pioneer in the detection of ARV in wild birds in Brazil, reporting for the first time the occurrence of PBV in wild species of bird *Guira guira*. Additional studies about epidemiological monitoring of infectious agents in wild birds are necessary, especially when involving segmented genome viruses, where processes of transmission, evolution and adaptation to new environments and hosts occur faster. Molecular characterization and phylogenetic analysis support to understand the epidemiology, origin, evolution and emergence of new viruses that may pose problems in the context of One Health.

## Author contributions

**Methodology**: Pereira, D., Souto, L.C.S., Guerra, S.F.S., Penha-Júnior, E.T., Lobo, P.S., Ramos, B.A., Chagas, L.L., Freitas, M.N.O., Furtado, E.C.S., Rodrigues, J.C.P. **Investigation:** Pereira, D., Souto, L.C.S., Guerra, S.F.S., Penha-Júnior, E.T., Lobo, P.S., Ramos, B.A., Chagas, L.L., Freitas, M.N.O., Furtado, E.C.S., Rodrigues, J.C.P. **Data analysis:** Pinheiro, H.H.C. **Data curation:** Pinheiro, H.H.C., Guimarães, R.J.P.S. **Writing - Original Draft:** Pereira, D. **Writing - Proofreading and Editing:** Mascarenhas, J.D.P., Chagas, E.H.N. **Supervision:** Mascarenhas, J.D.P., Guerra, S.F.S., Soares, L.S. **Project administration:** Mascarenhas, J.D.P., Martins, L.C., Casseb, A.R.

## Conflicts of interest

The authors declare no conflicts of interest.

## Funding information

Pereira, D., Souto, L.C.S., Chagas, E.H.N and Rodrigues, J.C.P are recipients of scholarships from the Coordination for the Improvement of Higher Education Personnel (CAPES); Ramos, B.A and Mascarenhas, J.D.P are recipients of grants from the National Council for Scientific and Technological Development (CNPq).

## Ethical statement

This study is linked to the research project entitled “Epidemiological monitoring of infectious agents in free-ranging birds (class of birds - Linnaeus 1758) and hematophagous arthropods (phylum Arthropoda - Latreille 1829) of the Federal Rural University of Amazon, campus Belém campus”, according to the ethical principles of animal experiments. The project was approved by the Ethics Committee on the Use of Animals of the Federal Rural University of Amazon (CEUA/UFRA), certificate number 025/18, and by Biodiversity Information and Authorization System (SISBIO), under opinion n° 63488 −1.

## Acknowledgment

The authors thank the technical support of Gastroenteric Viruses Laboratory (LGAST) on the Virology Section of Evandro Chagas Institute, Secretarial of Health Surveillance, and Ministry of Health. The technical team of the Section of Arbovirology and Hemorrhagic Fevers of Evandro Chagas Institute, and the Molecular Biology Laboratory of the Institute of Health and Animal Production of the Federal Rural University of Amazon.

## Notes

### Competing Interest Statement

The authors have declared no competing interest.

## REFERENCES

1. Santos GGC, Matuella GA, Coraiola AM, Silva LCS, Lange RR, Santin E. Doenças de aves selvagens diagnosticadas na Universidade Federal do Paraná (2003-2007). Pesq Vet Bras 2008;28:565–570.

2. Morais ABC. Ocorrência de patógenos de origem bacteriana e viral e marcadores de virulência de *Escherichia coli* e *Rhodococcus equi* isolados das fezes de aves silvestres de cativeiro da fauna brasileira. Dissertação (Mestrado em Medicina Veterinária) – Botucatu, Universidade Estadual Paulista, 2014. 81p.

3. Alfieri AF, Tamehiro CY, Alfieri AA. Virus entérico RNA fita dupla, segmentado, em aves: rotavirus, reovirus e picobirnavirus. Semina 2000;21:101–103.

4. Luz MA, Bezerra DA, Silva RR, Guerreiro AN, Seixas LS, Bastos RKG, Mascarenhas JDP, Moraes CCG, Souza NF, Meneses AMC. Rotavirus research in Amazon wild birds kept in captivity in the state of Pará, Brazil. Rev Bras Med Vet 2014;36:167–173.

5. Santos PMS, Silva SGN, Fonseca CF, Oliveira JB. Parasitos de aves e mamíferos silvestres em cativeiro no estado de Pernambuco. Pesq Vet Bras 2015;35:788–794.

6. Bezerra DAM, Silva RR, Kaiano JHL, Oliveira DS, Gabbay YB, Linhares AC, Mascarenhas JDP. Detection, epidemiology and characterization of VP6 and VP7 genes of group D rotavirus in broiler chickens. Avian Pathol 2014;43:238–243.

7. Lu H, Tang Y, Dunn PA, Wallner-Pendleton EA, Lin L, Knoll EA. Isolation and molecular characterization of newly emerging avian reovirus variants and novel strains in Pennsylvania, USA, 2011–2014. Sci Rep-UK 2015;5:14727.

8. Noh JY, Lee DH, Lim TH, Lee JH, Day JM, Song CS. Isolation and genomic characterization of a novel avian orthoreovirus strain in Korea, 2014. Arch Virol 2018;163:1307–1316.

9. El Taweel A, Kandeil A, Barakat A, Rabiee OA, Kayali G, Ali MA. Diversity of Astroviruses Circulating in Humans, Bats, and Wild Birds in Egypt. Viruses 2020;12:485.

10. Hassan MM, El-Zowalaty ME, Islam A, Khan SA, Rahman MK, Jarhult JD, Hoque MA. Prevalence and Diversity of Avian Influenza Virus Hemagglutinin Sero-Subtypes in Poultry and Wild Birds in Bangladesh. Vet Sci 2020;7:73.

11. Vidaña B, Busquets N, Napp S, Pérez-Ramírez E, Jiménez-Clavero MA, Johnson N. The Role of Birds of Prey in West Nile Virus Epidemiology. Vaccines 2020;8:550.

12. Duarte-Júnior JWB, Chagas EHN, Serra ACS, Souto LCS, Penha-Júnior ET, Bandeira RS, Guimãraes RJPS, Oliveira HGS, Sousa TKS, Lopes CTA, Domingues SFS, Pinheiro HHC, Malik YS, Salvarani FM, Mascarenhas JDP. Ocurrence of rotavirus and picobirnavirus in wild and exotic avian from amazon Forest. PLoS Negl Trop Dis 2021;15:e0008792.

13. Rahman MM, Talukder A, Chowdhury MMH, Talukder R, Akter R. Coronaviruses in wild birds – A potential and suitable vector for global distribution. Vet Med Sci 2021;7:264–272.

14. Jindal N, Patnayak DP, Chander Y, Ziegler AF, Goyal SM. Detection and molecular characterization of enteric viruses from poult enteritis syndrome in turkeys. Poultry Sci 2010;89:217–226.

15. Day JM, Zsak L. Molecular characterization of enteric picornaviruses in archived turkey and chicken samples from the United States. Avian Dis 2016;60:500–505.

16. Tang Y, Lin L, Sebastian A, Lu H. Detection and characterization of two co-infection variant strains of avian orthoreovirus (ARV) in young layer chickens using next-generation sequencing (NGS). Sci Rep-UK 2016;6:24519.

17. Pankovics P, Boros A, Nemes C, Kapusinszky B, Delwart E, Reuter, G. Molecular characterization of a novel picobirnavirus in a chicken. Arch Virol 2018;163:3455–3458.

18. Davis JF, Kulkarni A, Fletcher O. Reovirus infections in young broiler chickens. Avian Dis 2013;57:321–325.

19. Assunção TRS, Palka APG, Pavoni DP. Reovirose aviária: um panorama. Rev Educ Contin Med Vet Zootec CRMV-SP 2018;16:48–59.

20. Silva RR, Bezerra DAM, Kaiano JHL, Oliveira DS, Silvestre RVD, Gabbay YB, Ganesh B, Mascarenhas JDP. Genogroup I avian picobirnavirus detected in Brazilian broiler chickens: a molecular epidemiology study. J Gen Virol 2014;95:117–122.

21. Verma H, Mor SK, Erber J, Goyal SM. Prevalence and complete genome characterization of turkey picobirnaviruses. Infect Gen Evol 2015;30:134–139.

22. International Commitee on Taxonomy of Viruses (ICTV). ICTV Taxonomy history: *Orthoreovirus*. 2019. Disponível em: <https://talk.ictvonline.org/taxonomy/p/taxonomy-history?taxnode_id=201904969>. Acesso em: 09/10/2021.

23. Jones RC. Avian reovirus infections. Rev Sci Tech Oie 2000;19:614–619.

24. Dandár E, Bálint A, Kecskeméti S, Szentpáli-Gavallér K, Kisfali P, Melegh B, Farkas SL, Bányai K. Detection and characterization of a divergent avian reovirus strain from a broiler chicken with central nervous system disease. Arch Virol 2013;158:2583–2588.

25. International Commitee on Taxonomy of Viruses (ICTV). ICTV Taxonomy history: *Picobirnavirus*. 2019. Disponível em: <https://talk.ictvonline.org/taxonomy/p/taxonomy-history?taxnode_id=201904333>. Acesso em: 09/10/2020.

26. Ganesh B, Masachessi G, Mladenova Z. Animal picobirnavirus. Virusdisease 2014;25:223–238.

27. Nates SV, Gatti MSV, Ludert JE. The picobirnavirus: an integrated view on its biology, epidemiology and pathogenic potential. Future Virol 2011;6:223–235.

28. Gwynne JA, Ridgely RS, Argel M, Tudor G. Guia Aves do Brasil: Pantanal & Cerrado. São Paulo: Editora Horizonte, 2010, p. 336.

29. Comitê Brasileiro de Registros Ornitológicos (CBRO). Lista de aves do Brasil. 11° edição. 2014. Disponível em: <http://www.cbro.org.br>. Acesso em: 14/12/2021.

30. Sigrist T. Guia de Campo Avis Brasilis: Avifauna Brasileira. Pranchas, 2014.

31. Boom R, Sol CJA, Salimans MMM, Jansen CL, Wertheim-Van Dillen PME, Van Der Noordaa J. Rapid and simple method for purifications of nucleic acids. J Clin Microbiol 1990;28:495–503.

32. Pereira HG, Azevedo RS, Leite JP, Barth OM, Sutmoller F, Farias V, Vidal MN. Comparison of polyacrylamide gel electrophoresis (PAGE), immuno-electron microscopy (IME) and enzyme immunoassay (EIA) for the rapid diagnosis of rotavírus infection in children. Mem I Oswaldo Cruz 1983;78:483–490.

33. Zhang Y, Liu M, Shuidong O, Hu QL, Guo DC, Chen HY, Han Z. Detection and identification of avian, duck, and goose reoviruses by RT-PCR: goose and duck reoviruses are part of the same genogroup in the genus Orthoreovirus. Arch Virol 2006;151:1525–1538.

34. Rosen BI, Fang ZY, Glass RI, Monroe SS. Cloning of human picobirnavirus genomic segments and development of an RT-PCR detection assay. Virology 2000;277:316–329.

35. Malik YS, Sircar S, Dhama K, Singh R, Ghosh S, Bányai K, Vlasova AN, Nadia T, Singh RK. Molecular epidemiology and characterization of picobirnaviruses in small ruminant populations in India. Infect Genet Evol 2018;63:39–42.

36. Tamura K, Stecher G, Peterson D, Filipski A, Kumar S. MEGA6: molecular evolutionary genetics analysis version 6.0. Mol Biol Evol 2013;30:2725–2729.

37. Altschul SF, Gish W, Miller W, Myers EW, Lipman DJ. Basic local alignment search tool. J Mol Biol 1990;215:403–410.

38. Kimura M. A simple method for estimating evolutionary rates of base substitutions through comparative studies of nucleotide sequences. J Mol Evol 1980;16:111–120.

39. Felsenstein J. Confidence limits on phylogenies: An approach using the bootstrap. Evolution 1985;39:783–791.

40. Kearse M, Moir R, Wilson A, Stones-Havas S, Cheung M, Sturrock S, Thierer T. Geneious Basic: an integrated and extendable desktop software platform for the organization and analysis of sequence data. Bioinformatics 2012;28:1647–1649.

41. Ayres M, Ayres JRM, Ayres DL, Santos AAS. BioEstat: Aplicações estatísticas nas áreas das ciências biomédicas. 5^a^ Edição. 2007. Instituto de Desenvolvimento Sustentável Mamirauá, Belém, Pará, Brasil.

42. Wang H, Gao B, Chen H, Diao Y, Tang Y. Isolation and characterization of a variant duck orthoreovirus causing spleen necrosis in Peking ducks, China. Transbound Emerg Dis 2019;66:2033–2044.

43. Wong AH, Cheng PKC, Lai MYY, Leung PCK, Wong KKY, Lee WY, Lim WWL. Virulence potential of fusogenic orthoreoviruses. Emerg Infect Dis 2012;18:944.

44. Ogasawara Y, Ueda H, Kikuchi N, Kirisawa R. Isolation and genomic characterization of a novel orthoreovirus from a brown-eared bulbul *(Hypsipetes amaurotis)* in Japan. J Gen Virol 2015;96:1777–1786.

45. McDonald SM, Nelson MI, Turner PE, Patton JT. Reassortment in segmented RNA viruses: mechanisms and outcomes. Nat Ver Microbiol 2016;4:448–460.

46. De Sousa TN, Silva RVS, Evangelista BBC, Freire SM. Prevalência das zoonoses parasitárias e a sua relação com as aves silvestres no nordeste do Brasil. J Interdiscip Biociênc 2018;3:39–44.

47. Chagas EHN. Pesquisa de picobirnavírus e reovírus em espécimes fecais de animais das mesorregiões metropolitana de Belém e nordeste do estado do Pará no período de 2014 a 2016. Dissertação (Mestrado em Virologia) – Ananindeua, Instituto Evandro Chagas, 2018. 91p.

48. Guerreiro AN, Moraes CCG, Marinho ANR, Barros BCV, Bezerra DAM, Bandeira RS, Silva RR, Rocha DCC, Meneses AMC, Luz MA, Paz GS, Mascarenhas JDP. Investigation of enteric viruses in the feces of neotropical migratory birds captured on the coast of the State of Pará, Brazil. Braz J Poultry Sci 2018;20:161–168.

49. Yun T, Yu B, Ni Z, Ye W, Chen L, Hua J, Zhang C. Isolation and genomic characterization of a classical Muscovy duck reovirus isolated in Zhejiang, China. Infect Genet Evol 2013;20:444–453.

50. Masachessi G, Martinez LC, Giordano MO, Barril PA, Isa BM, Ferreyra L, Villareal D, Carello M, Asis C, Nates, S.V. Picobirnavirus (PBV) natural hosts in captivity and virus excretion pattern in infected animals. Arch Virol 2007;152:989–998.

51. Ribeiro AF, Silva RR, Bezerra DAM, Bandeira RS, Penha-Júnior ET, Castro CMO, Mascarenhas JDP. Picobirnavirus genogroup 2 in broiler chickens of the metropolitan Belém mesoregion–PA-Brazil. Braz J Anim Env Res 2019;2:2033–2050.

52. Silva RR. Detecção, epidemiologia e análise molecular de rotavirus, picobirnavirus e reovírus em aves de corte criadas em granjas na mesorregião metropolitana de Belém, Pará, Brasil. Tese (Doutorado em Doenças Tropicais) – Belém, Federal University of Pará, 2012. 153p.

53. Woo PCY, Teng JLL, Bai R, Tang Y, Wong AYP, Li KSM, Lam CSF, Fan RYY, Lau SKP, Yuen KY. Novel picobirnaviruses in respiratory and alimentary tracts of cattle and monkeys with large intra-and inter-host diversity. Viruses 2019;11:574.

54. Wille M, Eden JS, Shi M, Klaassen M, Hurt AC, Holmes EC. Virus–virus interactions and host ecology are associated with RNA virome structure in wild birds. Mol Ecol 2018;27:5263–5278.

55. Styś-Fijoł N, Kozdruń W, Czekaj H. Detection of avian reoviruses in wild birds in Poland. J Vet Res 2017;61: 239–245.

56. Al-Ebshahy E, Mohamed S, Abas O. First report of seroprevalence and genetic characterization of avian orthoreovirus in Egypt. Trop Anim Health Prod 2019;52:1049–1054.

57. Bányai K, Dandár E, Dorsey KM, Mató T, Palya V. The genomic constellation of a novel avian orthoreovirus strain associated with runting-stunting syndrome in broilers. Virus Genes 2011;42:82–89.

58. Reck C, Menin A, Pilati C, Miletti LC. Características clínicas e anatomo-histopatologicas da infecção experimental mista por Orthoreovirus aviario e Mycoplasma synoviae em frangos de corte. Pesqui Vet Brasil 2012;32:687–691.

59. Lima DA, Cibulski SP, Tochetto C, Varela APM, Finkler F, Teixeira TF, Loiko MR, Cerva C, Junqueira DM, Mayer FQ, Roehe PM. The intestinal virome of malabsorption syndrome-affected and unaffected broilers through shotgun metagenomics. Virus Res 2019;261:9–20.

60. Masachessi G, Martinez LC, Ganesh B, Giordano MO, Barril PA, Isa MB, Ibars A, Pavan JV, Nates SV. Establishment and maintenance of persistent infection by picobirnavirus in greater rhea *(Rhea Americana)*. Arch Virol 2012;157:2075–2082.

61. Navarro R, Chan Y, Nair R, Peda A, Aung MS, Ketzis J, Malik YS, Kobayashi N, Ghosh S. Molecular characterization of complete genomic segment-2 of picobirnavirus strains detected in a cat and a dog. Infect Genet Evol 2017;54:200–204.

62. Kumar N, Mascarenhas JDP, Ghosh S, Masachessi G, Bandeira RS, Nates SV, Dhama K, Singh RK, Malik YS. Picobirnavirus. In: Animal-Origin Viral Zoonoses. Y.S, Malik.; R.K, Singh.; K, Dhama. 1^a^ ed. Singapura: Springer Singapore, 2020, p. 291–312.

63. Ramesh A, Bailey ES, Ahyong V, Langelier C, Phelps M, Neff N, Sit R, Tato C, Derisi JL, Greer AG, Gray GC. Metagenomic characterization of swine slurry in a North American swine farm operation. Sci Rep 2021;11:16994.

64. Kashnikov AY, Epifanova NV, Novikova NA. Picobirnaviruses: prevalence, genetic diversity, detection methods. Vavilov Journal of Genetics and Breeding 2020;24:661–672.

